# Lowly polymorphic pathogen proteins may not be under evolutionary constraint

**DOI:** 10.64898/2026.10.14.759147

**Authors:** Ana Pinharanda, Sarah A Perkins, Angela M Early, Daniel E Neafsey

## Abstract

A central challenge in infectious disease biology is anticipating whether pathogen proteins targeted by vaccines, drugs, or diagnostics can evolve in ways that reduce intervention efficacy. Evolutionary constraint is therefore relevant to selecting targets likely to remain effective after deployment. Yet low diversity in contemporary pathogen populations is often interpreted as evidence of evolutionary constraint, an inference that can be misleading. In this Personal View, we use *Plasmodium falciparum* malaria to illustrate how contemporary population diversity and longer-term cross-species divergence can inform assessments of evolutionary constraint. A severe population bottleneck reduced diversity across the *P. falciparum* genome, meaning that low diversity in a particular protein-coding gene does not necessarily indicate unusual constraint. We place gene-level diversity in its genome-wide context, compare it with divergence across related ape-infecting malaria parasites, and examine how both relate to experimental essentiality. We then examine diversity and divergence along vaccine, drug, and diagnostic targets, considering them in relation to known resistance-associated mutations and other routes of evolutionary escape. We introduce PEEP (Protein Evolution Explorer for Plasmodium), a public interactive resource for exploring diversity and divergence across *P. falciparum* protein-coding genes and along individual proteins. Although our analyses focus on *P. falciparum* malaria, the implications extend across pathogens: low diversity alone is insufficient evidence of evolutionary constraint, and assessments of intervention durability should consider constraint alongside potential routes of evolutionary escape.

## Introduction

A central challenge in infectious disease biology is to anticipate how pathogen proteins targeted by an intervention will respond following implementation. For vaccines, drugs and diagnostics alike, the key question is whether the targeted protein can evolve in ways that compromise efficacy (1–3). In pathogen studies, proteins or genomic regions that harbor few polymorphisms are often described as conserved, with conservation viewed as a desirable property of intervention targets (4–10). However, a paucity of polymorphism is not sufficient to infer evolutionary constraint, nor to forecast the likelihood of resistance evolution.

The growing scale and accessibility of population genomic datasets for various pathogens makes this premise increasingly important. These datasets can be useful for disease surveillance because they show which variants are circulating and how allele frequencies change over time (11–18), but diversity reflects mutation, selection, genetic drift, recombination, demographic history, life history, and sampling bias. Low diversity therefore does not provide a direct measure of evolutionary constraint, especially in pathogens with low genome-wide diversity, including the causative agents of tuberculosis (*Mycobacterium tuberculosis*), leprosy (*Mycobacterium leprae*), bubonic plague (*Yersinia pestis*), and falciparum malaria (*Plasmodium falciparum*) (19–23).

Here we focus on *P. falciparum*, the deadliest human malaria parasite and the predominant cause of malaria in Africa, for which understanding evolutionary constraint has immediate translational relevance (**Figure 1A**). The evolutionary lineage leading to *P. falciparum* went through a severe bottleneck during a gorilla-to-human host switch (24), reducing diversity across the genome (**Figure 1B**) (24–26). In addition to this genome-wide demographic effect, selection can further shape variation at particular loci, with selective pressures differing across parasite life stages (27). A protein therefore may have little diversity within *P. falciparum*, without being unusually constrained. Analysis of protein sequence divergence among closely related ape-infecting malaria parasites, however, provides additional complementary information that strongly augments the application of diversity metrics for understanding constraint in *P. falciparum* proteins (**Box 1**).

**Figure 1.**
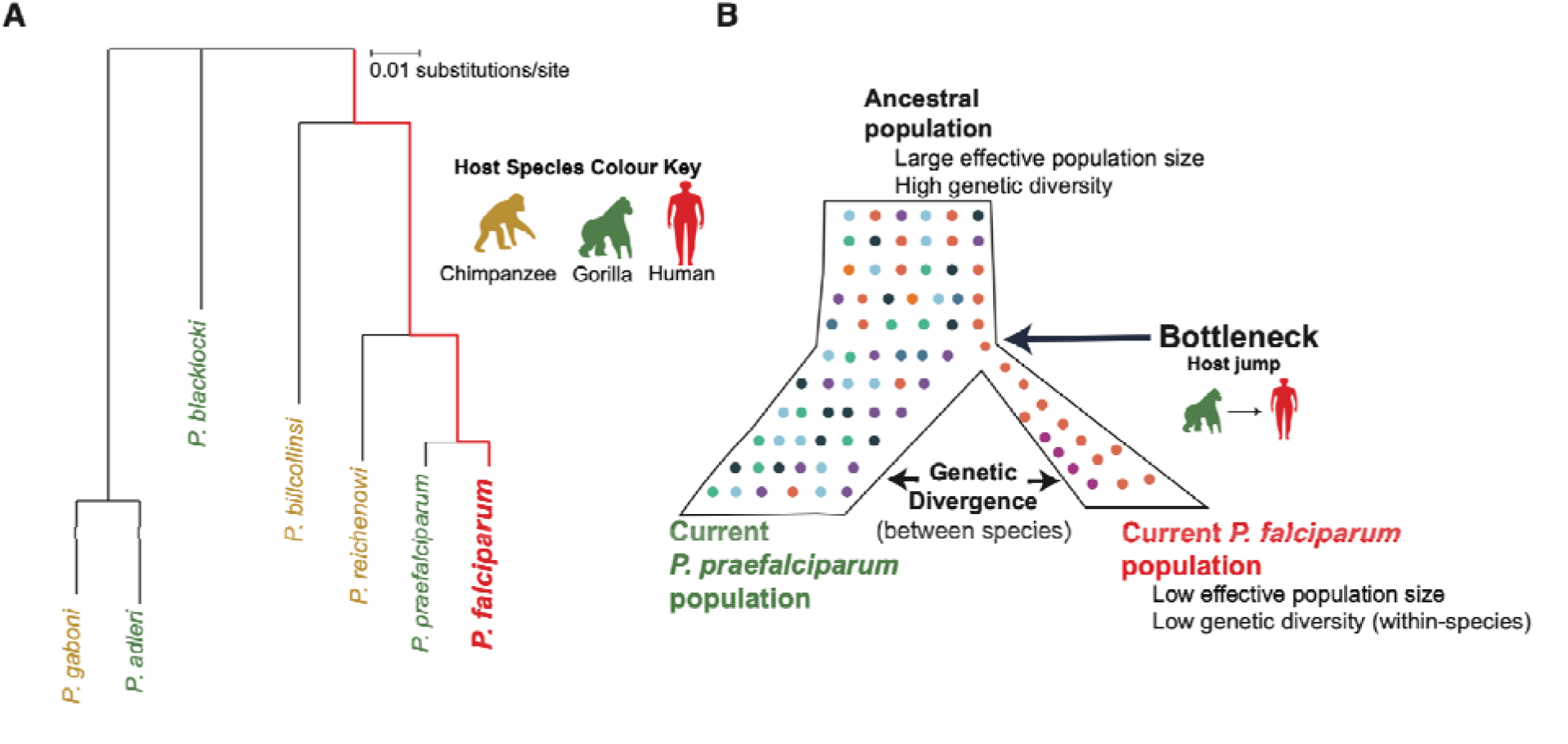
Evolutionary context for interpreting diversity and divergence in *P. falciparum*. **A) Species phylogeny of *P. falciparum* and its closest ape-infecting relatives inferred from single-copy orthologues.** The tree shows the relationships among the species included in the analysis, with *P. praefalciparum* as the sister lineage to *P. falciparum*. **B) Demographic bottleneck after the host jump into humans**. The lineage leading to *P. falciparum* underwent a severe reduction in effective population size, lowering diversity genome-wide.

For proteins targeted by antimalarial drugs, vaccines, or diagnostic tests, evolutionary constraint is important to understand the history of resistance emergence to diverse interventions. Mutations in Kelch13 (K13) confer partial resistance to artemisinin (28), a component of the global first-line drug treatment for falciparum malaria. Deletion of the *pfhrp2* gene, often together with its paralog *pfhrp3*, can cause failure of rapid diagnostic tests that target the PfHRP2 protein (29–31). Antigenic variation in the circumsporozoite protein (CSP) can affect vaccine efficacy (32), inspiring *pfcsp* surveillance efforts (33). Newer interventions are being developed under the specter of historical resistance emergence, including ganaplacide-lumefantrine combination therapy, a recently approved alternative to artemisinin-based combination therapy (ACTs) for which reduced susceptibility has been linked to mutations in the cyclic amine resistance locus (CARL) (34); rapid diagnostic tests that detect lactate dehydrogenase (PfLDH) (35–37); and the blood-stage vaccine candidate targeting reticulocyte-binding protein homologue 5 (RH5) (38). We use these examples to examine how evidence from individual polymorphisms, population-level diversity, and inter-species divergence should be interpreted when assessing evolutionary constraint and intervention durability, in the context of each target and the mechanism by which the intervention can fail.

### Box 1. Key concepts for interpreting diversity, divergence and constraint

**Sequence-based evidence**

Polymorphism A site or allele that differs among sampled sequences from the same population or species. A polymorphic site may be rare or common.

Diversity A quantitative measure of sequence variation among sampled sequences from the same population or species. Diversity reflects both the amount of variation and the frequencies of alleles. A variant found at high frequency contributes more to diversity than a singleton. Low diversity can reflect demography, evolutionary constraint, or recent positive selection that has reduced variation around a favored allele. Sampling and limited power to detect rare variants can also affect estimates of diversity.

Divergence A quantitative measure of sequence differences that have accumulated between species or lineages over evolutionary time. Divergence provides a longer-term view of evolutionary change. Factors influencing divergence include evolutionary constraint, random genetic drift, adaptation, and positive selection.

Conservation Sequence similarity within or across species. Conservation can be measured using percent identity or the amount of variation at individual sequence positions. It is often used as evidence of evolutionary constraint. However, conservation is the observed pattern, whereas constraint is the underlying evolutionary property inferred from that pattern.

**Evolutionary and demographic properties and processes**

Evolutionary constraint The degree to which amino-acid changes in a protein (or protein region) are selected against because they affect protein function or fitness. Reduced diversity and divergence can therefore be signals of stronger constraint, but constraint does not always result in conservation. For example, in populations with small or reduced effective population size (*Ne*), genetic drift can allow changes to persist or become fixed even when they are selected against.

Bottleneck A demographic reduction in population size that lowers genetic diversity across the genome. In *P. falciparum*, the bottleneck associated with the host jump into humans reduced diversity across the genome.

Effective population size (*Ne*) A measure of population size that reflects the strength of genetic drift. Lower Ne increases drift and reduces the efficiency of selection.

Anthropogenic selection Selection imposed by human activities or interventions. This can include selection from drugs, vaccines, diagnostics, vector control, or other public-health interventions.

**Target-assessment claims and outcomes**

Essentiality The extent to which an organism depends on a gene or protein for growth and survival under a particular experimental or biological condition. Essentiality screens estimate the effect of disrupting or depleting that gene product.

Durability The expected ability of an intervention target to remain effective after deployment.

Evolutionary escape Pathogen evolution that reduces the effectiveness of an intervention. The underlying genetic changes can include sequence variants, copy-number changes, gene deletion, repeat variation, or altered expression.

### Genome-wide context: diversity and divergence

To examine what divergence and diversity can tell us about evolutionary constraint in *P. falciparum*, we profile their genome-wide distributions across protein-coding genes (**Figure 2**), using divergence across related ape-infecting parasites and diversity within a contemporary *P. falciparum* population. Across the seven genomes analysed, 36,592 predicted protein-coding genes were assigned to 5,079 orthogroups, representing 99.5% of predicted protein-coding genes. Of these, 4,300 orthogroups contained sequences from all seven species, and 4,077 had a single sequence from each species (**Box 2**). This covers around 80% of the core nuclear *P. falciparum* genome which can be reliably analysed with short-read sequencing data (39).

**Figure 2.**
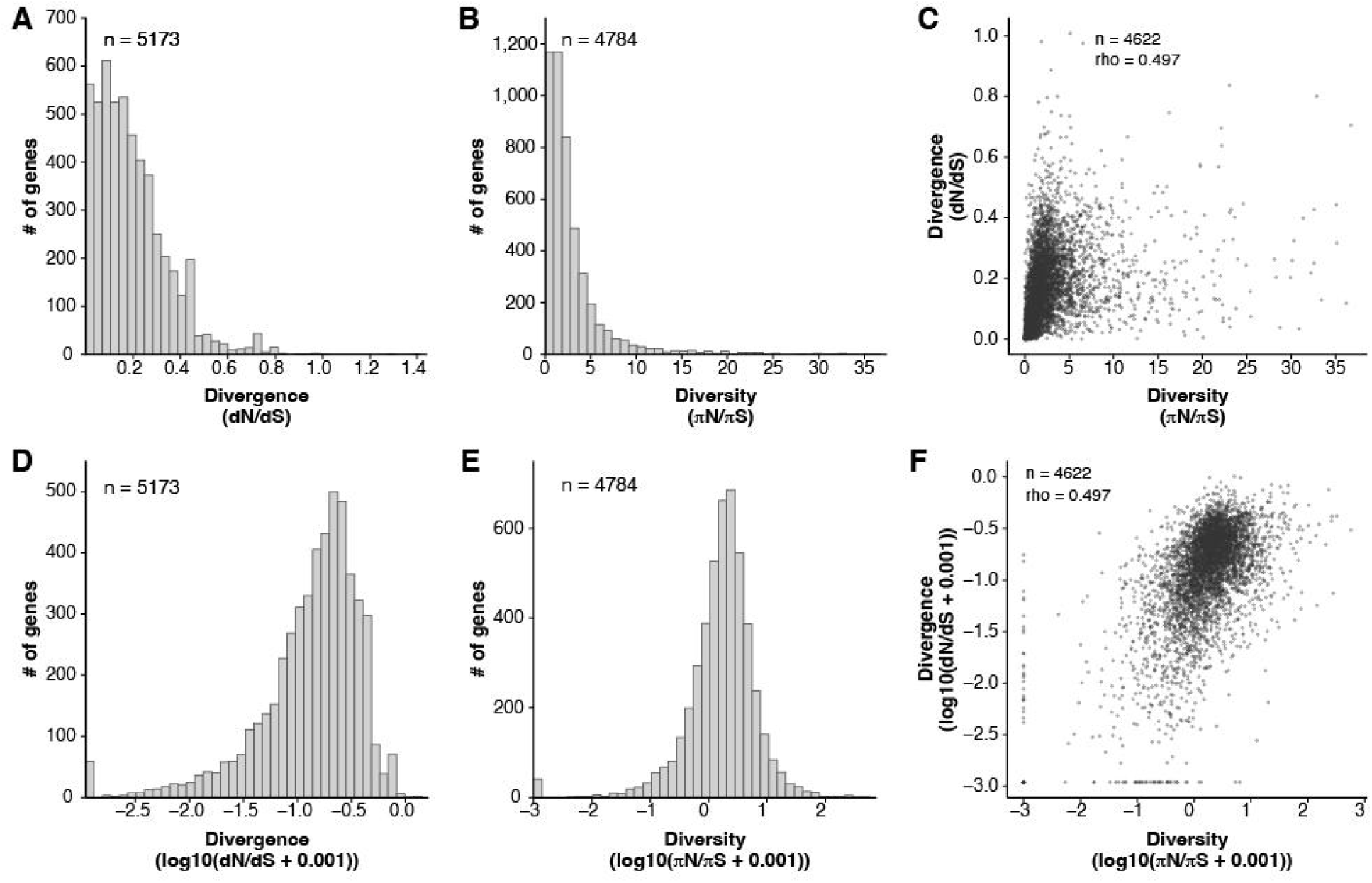
Genome-wide distributions of diversity and divergence across protein-coding genes. **A–C)** Gene-level divergence, diversity, and their relationship on the untransformed scale. **A)** Distribution of divergence (dN/dS) estimates across protein-coding orthogroups from closely related ape-infecting parasites. **B)** Distribution of diversity (πN/πS) estimates across *P. falciparum* protein-coding genes. Because of a small but long upper tail in the πN/πS distribution, the lower 99% is displayed (4,736 of 4,784 protein-coding genes); the upper 1% (48 genes; πN/πS > 36.7) falls outside the displayed range. **C)** Relationship between diversity and divergence for the 4,622 protein-coding genes with both estimates. The same πN/πS display threshold as in **B** is applied. **D–F)** The corresponding genome-wide distributions and relationship after log10 transformation, with all available protein-coding genes shown: **D)** divergence, **E)** diversity, and **F)** their relationship. Histogram bars show the number of protein-coding genes per bin in A, B, D, and E; points in C and F represent individual protein-coding genes. Divergence and diversity are moderately correlated (Spearman ρ = 0.497). Log transformation resolves relative differences among protein-coding genes across the skewed genome-wide distributions and is used for subsequent genome-wide figures.

The untransformed distributions illustrate an important limitation of diversity for inferring evolutionary constraint: many protein-coding genes have very little observed variation, providing limited resolution among genes at the low end of the distribution. At the same time, πN/πS is strongly right-skewed, with a long upper tail partly driven by genes with very low synonymous diversity (πS, **Box 2**), for which even modest nonsynonymous diversity can produce a large ratio (**Figure 2A–C**). Log transformation spreads the densely occupied portion of the distribution and compresses its long upper tail, making the relative positions of protein-coding genes within the genome-wide distribution easier to visualise (**Figure 2D–F**). We therefore use genome-wide percentile rank in subsequent comparisons (**Box 2)**, asking both whether a protein is unusually low in diversity and how its diversity compares with its divergence.

Divergence and diversity are moderately correlated across genes (Spearman ρ = 0.497), as expected because constraints on protein function and fitness can affect both nonsynonymous substitution between species and nonsynonymous polymorphism within *P. falciparum* (40). However, divergence varies considerably among genes with similarly low diversity (**Figure 2C, F**). For example, among the 4,622 protein-coding genes with both estimates, the 463 in the lowest 10th percentile for diversity (πN/πS ≤ 0.390) range from the 0.6th to the 97.4th percentile for divergence. Thus, even among proteins with similarly low contemporary diversity, some show almost no amino-acid divergence across related species, whereas others are among the most divergent proteins in the genome.

#### Box 2. Comparative metrics used in this analysis

**Orthogroups** are sets of protein-coding genes from different species descended from the same ancestral protein-coding gene. They allow diversity within a species to be compared with divergence across related species.

**π** measures nucleotide diversity as the average number of pairwise nucleotide differences per site among sampled sequences, accounting for their frequencies in the sample. **πN** is the average number of nonsynonymous nucleotide differences per nonsynonymous site, whereas **πS** is the average number of synonymous nucleotide differences per synonymous site. The ratio **πN/πS** summarizes nonsynonymous diversity relative to synonymous diversity.

**dN** estimates the rate of nonsynonymous substitutions per nonsynonymous site between species, capturing protein-changing divergence over evolutionary time. **dS** estimates synonymous substitutions per synonymous site over the same comparisons. Their ratio, **dN/dS**, or **ω**, scales nonsynonymous divergence by synonymous divergence. Low **ω** is consistent with stronger evolutionary constraint. Higher **ω** indicates greater relative nonsynonymous divergence, which can reflect relaxed constraint, drift, lineage-specific adaptation, positive selection, or technical factors such as alignment uncertainty.

**Codon-level πN** maps nonsynonymous diversity along a protein, while **codon-level dN** maps nonsynonymous divergence across related species. These profiles are useful because interventions often act on specific protein regions or sites: diagnostic antibodies bind epitopes, vaccine responses target epitopes or receptor-binding interfaces, and resistance-associated mutations occur at particular positions.

**Percentile rank** shows where each protein-coding gene falls within the genome-wide distribution for a given metric. We use mid-rank percentiles, scaled from 0 to 100, so genes with identical values, including zeros, are treated as tied rather than ordered arbitrarily. Higher percentiles indicate higher values relative to other genes.

### Beyond essentiality: introducing the Protein Evolution Explorer for *Plasmodium* (PEEP)

Deciding which parasite genes or proteins should be prioritised for further study or intervention development requires functional and translational evidence, including expression, accessibility, functional importance, experimental tractability, and intervention-specific phenotypes (41). Essentiality screens have long been used to aid drug and vaccine target prioritisation, but advances in high-throughput genetic perturbation techniques now allow essentiality to be assessed systematically across thousands of genes in an increasing range of organisms (42– 46). A recent piggyBac transposon mutagenesis screen for a zoonotic malaria parasite (*Plasmodium knowlesi*) provides the most comprehensive genome-wide assessment of blood-stage gene essentiality currently available for a malaria parasite, classifying genes as essential, intermediate, or dispensable from the density and distribution of tolerated insertions (46). We mapped these classes to *P. falciparum* through single-copy orthologues and examined how divergence and diversity were distributed across the three essentiality classes. For both, the distributions overlap extensively across classes (**Figure 3A and 3B**). Although median divergence and diversity are lower in essential genes, the differences are modest relative to the variance of the distributions. These classes therefore provide little basis for inferring the evolvability of a protein. Essentiality measures the effect of gene disruption on growth or survival under the conditions tested; it is not a measure of evolutionary constraint (44,47).

**Figure 3.**
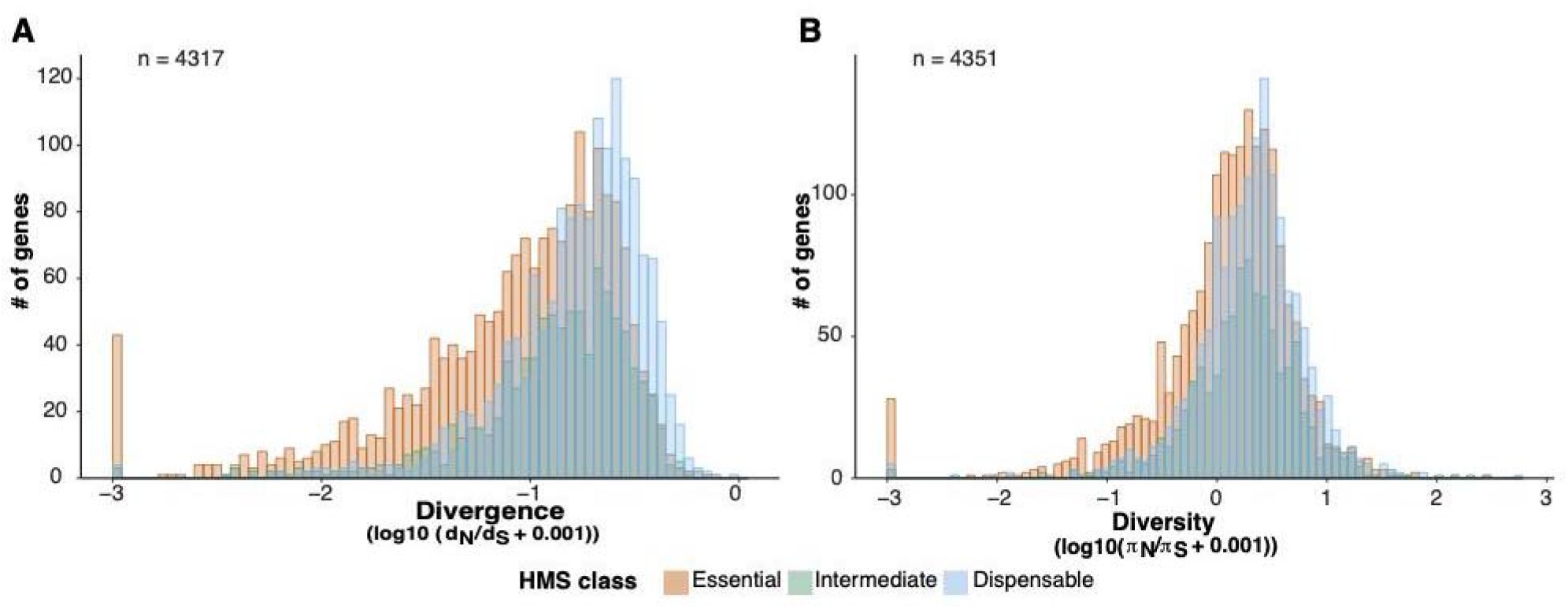
Relationship between diversity, divergence, and experimental essentiality across orthogroups. Distributions of gene-level **A)** divergence and **B)** diversity are shown across essential, intermediate, and dispensable HMS classes from the saturation-mutagenesis screen. Both metrics increased across HMS classes, with median diversity of 0.19, 0.265 and 0.338; and a median divergence of −0.954, −0.819 and −0.706, respectively, on the plotted log10 scale.

Unlike an essentiality screen performed under defined experimental conditions, diversity and divergence reflect evolutionary outcomes accumulated across populations, generations, and biological contexts. Genome-wide approaches are increasingly incorporating polymorphism alongside other genomic and functional features to prioritise candidate targets across the *P. falciparum* proteome (11,12). We developed the Protein Evolution Explorer for *Plasmodium* (PEEP) (https://laverania.shinyapps.io/peep/), an accompanying resource for this Personal View, to add divergence across related parasites to this genome-wide context and allow diversity and divergence to be examined at both gene and protein level. The examples that follow use PEEP to show how these comparisons inform different questions: conservation in vaccine targets, the location of resistance-associated mutations in drug targets, and routes of evolutionary escape from diagnostic detection.

### Vaccine targets: constraint in a genome-wide context

Malaria vaccine efficacy depends on immune recognition of parasite proteins, so amino-acid polymorphism in those proteins can affect how well vaccine-induced responses recognize circulating parasites. CSP is the circumsporozoite protein targeted by RTS,S/AS01 and R21/Matrix-M, the two malaria vaccines recommended by WHO for use in children in malaria-endemic areas (48,49). RH5 is involved in erythrocyte invasion, and is the target of a leading candidate blood-stage vaccine RH5.1/Matrix-M, which has progressed through phase 2b clinical testing (50).

Our analysis of CSP and RH5 exposes different problems in how vaccine target constraint assessment is inferred from genomic data. At a gene level, CSP ranks in the 46th percentile for divergence genome-wide, whereas RH5 ranks in the 91st percentile (**Figure 4A, B**). Both rank near the top of the genome-wide diversity distribution: CSP at the 97.6th percentile and RH5 at the 98th percentile (**Figure 4C, D**). RH5 therefore ranks high for both diversity and divergence relative to other genes, despite its longstanding description as conserved (10,51–71).

**Figure 4.**
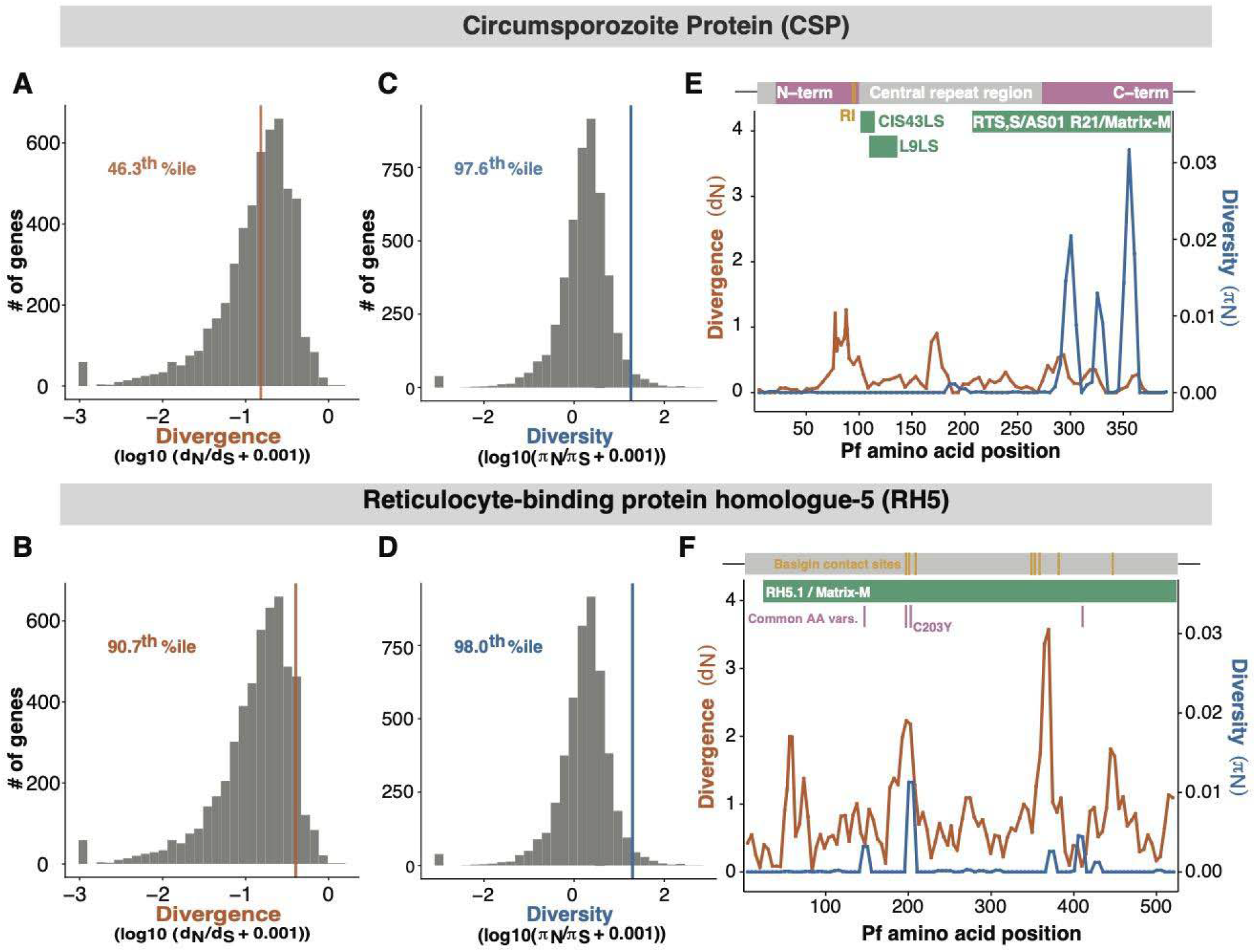
Diversity and divergence in CSP and RH5. **A, B)** Genome-wide distributions of protein-coding gene-level divergence, with CSP and RH5 marked by orange lines. **C, D)** Genome-wide distributions of protein-coding gene-level diversity, with CSP and RH5 marked by blue lines. Percentile ranks indicate the position of each gene within the corresponding genome-wide distribution. **E, F)** CSP and RH5 protein-level profiles showing codon-level nonsynonymous diversity and divergence, mapped to *P. falciparum* protein coordinates with functional and intervention-relevant annotations. CSP annotations include the N-terminal, central repeat and C-terminal regions, vaccine-targeted regions, and monoclonal-antibody binding regions. RH5 annotations include the RH5.1/Matrix-M vaccine construct, basigin-contact sites, and commonly reported amino-acid variants. C203Y is highlighted because it is a commonly reported PfRH5 polymorphism.

At the protein level, CSP shows that regions with modest divergence can still have appreciable diversity, particularly in the setting of strong immune selection. Meanwhile, RH5 shows substantial divergence across species, including near sites involved in host receptor binding encompassed by the vaccine construct (**Figure 4E, 4F**). RTS,S/AS01 and R21/Matrix-M include part of the central repeat region and the C-terminal region (C-term) of CSP, while candidate monoclonal antibodies (mAbs) target epitopes toward the N-terminal (N-term) side of the central repeat. Specifically, the CIS43LS mAb targets the junctional epitope between the N-term and central repeat region (72,73), whereas the L9LS mAb targets the minor repeats at the N-term of the central repeat region (74,75). The C-term shows high polymorphism (76), consistent with concerns about allele-specific protection (32), whereas the N-term has been described often as relatively conserved (77–81). Yet the N-term shows appreciable divergence across related parasites despite lower diversity within *P. falciparum* (**Figure 4E**), suggesting that mAbs or vaccines targeting the N-term may not be immune to an evolutionary escape response. The central repeat region is difficult to align and should be interpreted cautiously.

RH5 is an example of how observations of few polymorphisms can, without appropriate contextualization, gradually build into broader claims of conservation (10,51–62,65). An early, frequently cited study by Hayton and colleagues identified 10 polymorphic amino-acid positions across 18 *P. falciparum* laboratory strains (64). One year later, Baum and colleagues identified six nonsynonymous (and no synonymous) polymorphisms across seven laboratory strains and, notably, described this pattern as “suggestive, though not significantly so, of selection favouring diversity in the PfRh5 protein” (63). By 2011, Douglas and colleagues described the 10 sites reported by Hayton as “limited polymorphism” (66). In 2013, analysis of a much larger panel of 227 MalariaGEN samples by Bustamante and colleagues identified 12 nonsynonymous polymorphisms (67). Later that year, a study citing this analysis referred to PfRH5 as having “very few coding polymorphisms” and as a “relatively conserved merozoite surface antigen” (68). Subsequent work cited these polymorphism studies alongside evidence of the essential role of RH5 in erythrocyte invasion, and described RH5 more broadly as a “conserved parasite ligand” (69). By 2018, Ouattara and colleagues cited Hayton *et al*. 2008 (64) and Bustamante *et al*. 2013 (67) as evidence that *pfrh5* exhibits “limited polymorphism” and suggested that its functional requirement for erythrocyte invasion would “exert strong purifying selection on the locus and might constrain the genetic diversity of *pfrh5*” (10). Similar descriptions persist in recent literature, where RH5 continues to be described as a “conserved” or “highly-conserved” protein (70,71).

Those early observations established that relatively few polymorphisms had been detected in those samples; the number of observed polymorphisms alone, however, does not establish that RH5 is unusually low in polymorphism genome-wide or that it is evolutionarily constrained. Interpreting such a count requires both a denominator - the number of sites in a protein - and genome-wide context from other proteins analysed in the same population or species. Indeed, RH5 has 25 codons with detectable nonsynonymous polymorphism in the West African dataset analysed here, placing it near the middle of the genome-wide distribution by number of nonsynonymous-polymorphic codons. This is consistent with other genome-wide and expanded population analyses. For example, Chou and colleagues placed RH5 at approximately the 52nd percentile for SNPs per Kb across the *P. falciparum* proteome (12), while Mangou and colleagues identified substantially more RH5 polymorphisms than earlier studies (55).

The diversity and divergence analyses show that the early polymorphism counts provide an incomplete picture of RH5 evolution. In contemporary *P. falciparum*, RH5 is not unusually low in diversity relative to other proteins, while across related species it has accumulated substantial amino-acid divergence, including within the structural core and near the basigin-binding interface involved in red-blood-cell invasion (**Figure 4F**) (26,82). Taken together, these patterns provide little evidence that RH5 is unusually constrained on the basis of sequence conservation alone.

This does not undermine RH5 as a promising vaccine target or predict a route of vaccine escape. However, if RH5-based vaccines move towards wider use, its longstanding description as conserved should not by itself justify expectations of greater durability, nor warrant less intensive surveillance than what is currently being conducted for other more polymorphic vaccine targets such as CSP.

### Drug resistance-associated proteins: selected mutations and regional divergence

Drug treatment can drive resistance-associated mutations to higher frequency by favoring parasites that survive under drug pressure, even when those mutations carry fitness costs in the absence of treatment (83). Mutations in K13 are the main genetic determinants associated with reduced susceptibility to artemisinin, a component of first-line treatment for *P. falciparum* malaria worldwide (84–86). Ganaplacide-lumefantrine is advancing as a non-artemisinin antimalarial combination (87), and reduced susceptibility to ganaplacide has been linked to variation in the cyclic amine resistance locus, CARL (88,89). The diversity profiles analysed here are from West African parasites, where artemisinin resistance has not substantially shaped polymorphism in the K13 propeller domain (84).

Our analysis of K13 and CARL asks whether resistance-associated polymorphisms occur in parts of these proteins where divergent amino-acid substitutions have also accumulated across related parasites. At the protein-coding gene level, K13 ranks at the 8^th^ percentile for divergence genome-wide, whereas CARL ranks at the 63^rd^ percentile (**Figure 5A, 5B**). Their diversity rankings differ in the opposite direction: K13 ranks at the 85th percentile, compared with the 40th percentile for CARL (**Figure 5C, 5D**).

**Figure 5.**
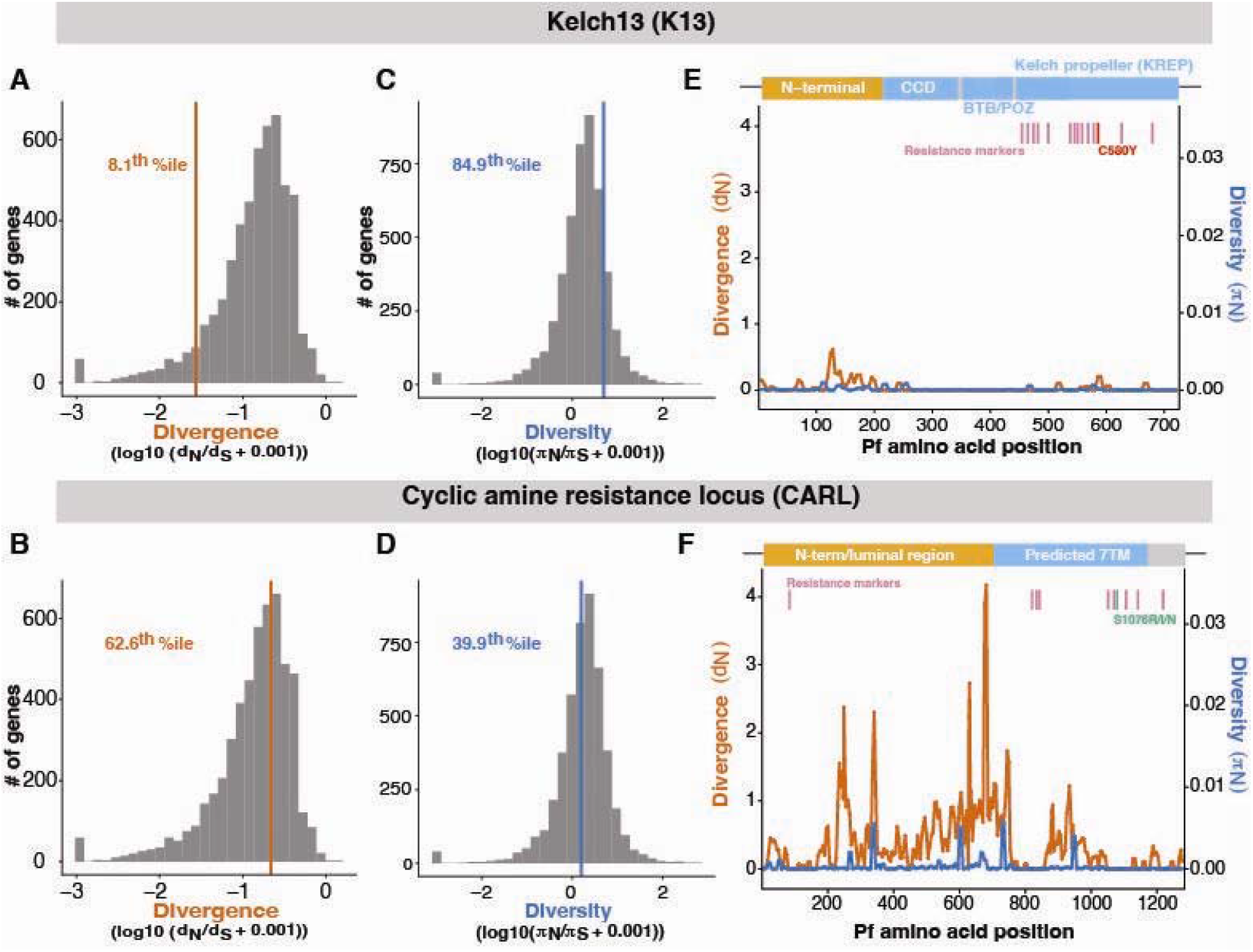
Diversity and divergence in K13 and CARL. **A, B)** Genome-wide distributions of protein-coding gene-level divergence, with K13 and CARL marked by orange lines. **C, D)** Genome-wide distributions of protein-coding gene-level diversity, with K13 and CARL marked by blue lines. Percentile ranks indicate the position of each gene within the corresponding genome-wide distribution. **E, F)** K13 and CARL protein-level profiles showing codon-level nonsynonymous diversity and divergence, mapped to *P. falciparum* protein coordinates. K13 annotations include the N-terminal region, CCD, BTB/POZ and Kelch propeller domains, together with validated resistance markers; C580Y is highlighted as a well-established artemisinin-resistance mutation. CARL annotations include the N-terminal/luminal and predicted transmembrane regions and laboratory-selected resistance substitutions; S1076R/I/N is highlighted because substitutions at this position produced the largest reported reduction in susceptibility in vitro.

The amino-acid level profiles show that the relationship between resistance-associated sites and divergence differs markedly between the two proteins (**Figure 5E, 5F**). In K13, resistance-associated mutations are concentrated in the propeller domain. The validated resistance sites are not themselves divergence peaks; for example, divergence is near zero at C469, R561, and C580 (**Figure 5E**). However, resistance sites occur close to positions with elevated divergence within the propeller domain. The diversity profile shown here is estimated from West African samples, where C580Y remains rare and partial resistance has more often been associated with other K13 mutations, including R561H and C469Y/F (90). By contrast, C580Y reached high frequency in Southeast Asian populations under artemisinin selection (91), illustrating how recent drug selection can drive variants to high frequencies at sites with little divergence. The Southeast Asian pattern therefore reflects recent drug-mediated selection, whereas the West African profile more closely reflects the other evolutionary forces shaping K13 diversity. The cross-species comparison, however, shows that the propeller domain had accumulated amino-acid substitutions over deeper evolutionary time. Thus, even when little polymorphism was observed before widespread resistance, divergence showed that amino-acid evolution had occurred in the propeller domain, providing evidence of evolutionary change that was not captured by diversity alone.

CARL shows the converse pattern. The protein contains several regions of elevated divergence, including residues in the predicted membrane-associated portion, but the annotated laboratory-selected resistance substitutions do not coincide with these regions (**Figure 5F**). For example, S1076N produced the largest reported reduction in susceptibility in vitro to GNF179, an experimental antimalarial closely related to ganaplacide (89), yet position 1076 shows no detectable diversity or divergence in our dataset. The same pattern extends across to the other annotated laboratory selection substitutions: they occur at sites with little or no detectable divergence and diversity, despite substantial divergence elsewhere in CARL.

The contrast between K13 and CARL suggests that resistance-associated mutations can arise in regions with very different histories of amino-acid change. For CARL, the absence of divergence at the laboratory-selected resistance sites is potentially encouraging: although these substitutions can reduce susceptibility in vitro, there is little evidence from related parasites that amino-acid substitutions have accumulated at the same sites. Whether these substitutions can arise and increase in frequency under ganaplacide selection in natural populations remains to be tested.

### Diagnostic antigens: different routes to escape

Antigen-based rapid diagnostic tests depend on parasite proteins being present and recognised by diagnostic antibodies (29,92,93). LDH-based tests are generally less sensitive than HRP2-based tests when HRP2 is present, but provide an alternative when HRP2-based detection is compromised and are already used alone or in combination with HRP2-based detection (37,94,95).

HRP2 and LDH illustrate how the relevance of diversity and divergence depends on the route by which diagnostic detection can fail. At the gene level, HRP2 ranks at the 76th percentile for divergence genome-wide, whereas LDH ranks at the 9th percentile (**Figure 6A, B**). The HRP2 estimate should be interpreted cautiously because its orthogroup contains both HRP2-like and HRP3-like sequences and the repeat-rich regions align poorly across species. It therefore provides broad evidence of divergence in this histidine-rich protein family rather than a precise estimate for HRP2. LDH, by contrast, has low divergence relative to other genes. For diversity, LDH ranks at the 96th percentile genome-wide (**Figure 6C**); a gene-level diversity estimate could not be calculated for HRP2 due to poor short-read sequence alignment quality.

**Figure 6.**
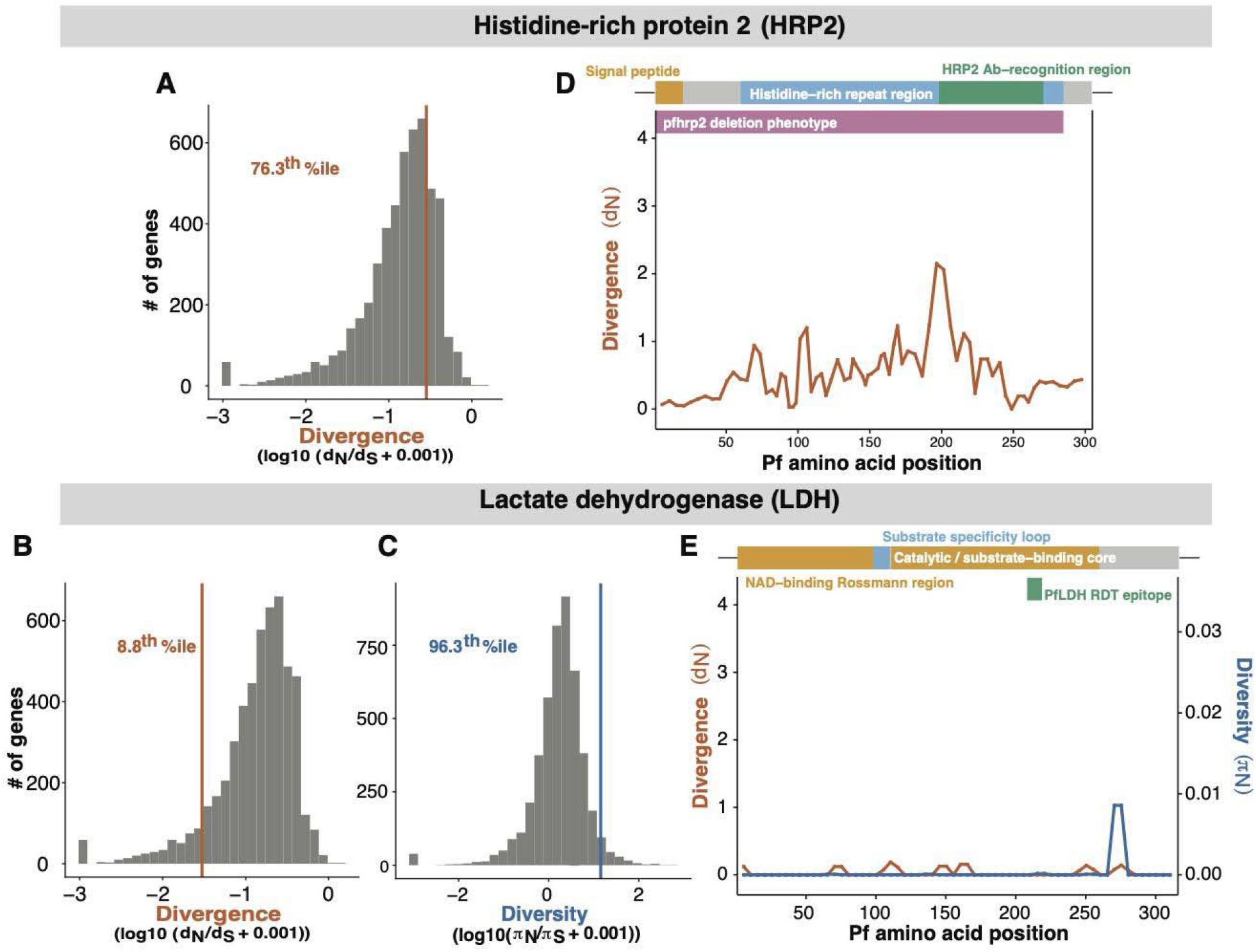
Diversity and divergence in HRP2 and LDH. **A, B**) Genome-wide distributions of protein-coding gene-level divergence, with HRP2 and LDH marked by orange lines. **C**) Genome-wide distribution of protein-coding gene-level diversity for LDH marked by a blue line; HRP2 is not shown because the gene-level diversity ratio could not be calculated. Percentile ranks indicate the position of each gene within the corresponding genome-wide distribution. **D, E**) HRP2 and LDH protein-level profiles showing codon-level nonsynonymous diversity and divergence, mapped to *P. falciparum* protein coordinates. HRP2 annotations include the signal peptide, histidine-rich repeat region, antibody-recognition region, and *pfhrp2* deletion phenotype. LDH annotations include the NAD-binding Rossmann region, substrate-specificity loop, catalytic/substrate-binding core, and a known RDT epitope spanning residues 90–210. The HRP2 divergence profile should be interpreted cautiously because its orthogroup contains HRP2-like and HRP3-like sequences and the repeat-rich regions align poorly across species.

Complete *pfhrp2* deletion—the major established route of diagnostic escape—is not represented by either profile. For LDH, the annotated RDT epitope spans residues 90–210, across which most positions show no detectable diversity or divergence despite the high gene-level diversity ranking (**Figure 6D, 6E**).

The purpose of this vignette is not to rank HRP2 and LDH as diagnostic targets, but to show that diversity and divergence may not always be informative. For HRP2, deletion and repeat variation are central and are not captured by these measures. For LDH, diversity and divergence can be interpreted more directly in relation to the region recognised by diagnostic antibodies; and the limited diversity and divergence across most of this region is encouraging for the durability of LDH-based detection against escape through amino-acid change.

### From evolutionary constraint to intervention durability

Pathogen intervention targets are often described as evolutionarily constrained on the basis of limited polymorphism or evidence of essentiality, without considering the additional genomic data that could support or challenge that inference. This matters because claims of constraint can shape target prioritisation and the focus of genomic surveillance after deployment. The examples considered here show that interpreting diversity and divergence requires considering population history and genome-wide context, the scale at which they are examined—from the full-length protein to individual regions and amino-acid positions—and the mechanism by which the intervention can fail. PEEP brings population diversity and divergence across related parasites together for *P. falciparum*, allowing both to be examined in the same genome-wide and protein-level framework. Users can add their own coordinates and annotations, allowing the profiles to be explored alongside features relevant to their question. Similar approaches could be developed for other pathogens as genomic resources expand.

The increasing availability of population, comparative, functional, and experimental data creates an opportunity to eventually move from assessing evolutionary constraint towards predicting evolutionary escape. In viral systems, EVEscape provides an example of what this might look like, using a deep-learning framework that combines information learned from sequence data with structural and biophysical information to predict antibody escape (96). Comparable approaches remain more difficult for bacterial and eukaryotic pathogens, where data linking genotype to intervention-relevant phenotypes are more limited. The longer-term opportunity is to bring population variation and cross-species divergence together with experimental and intervention-specific data, and ask whether these can be used to predict evolutionary escape as new interventions are deployed.

## Contributors

D.E.N. conceptualised the study; A.P., S.A.P., A.M.E., and D.E.N. developed the methodology; A.P. and S.A.P. performed the formal analyses and curated the data; A.P. developed the software and created the visualisations; A.P. wrote the original draft; A.P., S.A.P., A.M.E., and D.E.N. reviewed and edited the manuscript; A.M.E. and D.E.N. supervised the work.

## Acknowledgments

We thank Melodi Anahtar for comments on and careful reading of the manuscript, and Jacob A Tenessen and Peter H Culviner for helpful discussions. This work was funded in whole or in part with Federal funds from the National Institute of Allergy and Infectious Diseases, National Institutes of Health, Department of Health and Human Services, under Award Number U19AI110818. S.A.P. was supported by the National Institute of Allergy and Infectious Diseases of the National Institutes of Health under Award Number F31AI194773. The funders had no role in the preparation of the manuscript or the decision to submit it for publication.

## Data and code availability

The Protein Evolution Explorer for *Plasmodium* (PEEP) is publicly accessible at https://laverania.shinyapps.io/peep/. The code used for the analyses and generation of figures in this study, together with the code and precomputed data used to build and deploy PEEP, is publicly available at https://github.com/apinharanda/laverania.git. The repository also documents the analysis workflow.

## AI declaration

OpenAI ChatGPT (GPT-4.1 and GPT-5.6 Sol) was used during development of the Protein Evolution Explorer for *Plasmodium* (PEEP) to assist with integrating existing author-developed analysis code into the Shiny application and with modifying, writing, and troubleshooting code for the user interface and application functionality. The underlying analytical approaches and analysis code were developed by the authors independently of ChatGPT. The authors defined the functionality of PEEP, reviewed the AI-assisted code, tested and verified the application and its outputs. ChatGPT was also used during manuscript preparation for spelling, grammar, and language editing. All AI-assisted material was reviewed by the authors.

